# Metabolic profiling stratifies colorectal cancer and reveals adenosylhomocysteinase as a therapeutic target

**DOI:** 10.1101/2023.03.12.531945

**Authors:** Johan Vande Voorde, Arafath K. Najumudeen, Rory T. Steven, Chelsea J. Nikula, Alex Dexter, Lucas B. Zeiger, Efstathios A. Elia, Ammar Nasif, Ariadna Gonzalez-Fernandez, Teresa Murta, Michael Gillespie, Catriona A. Ford, Tamsin R.M. Lannagan, Nikola Vlahov, Rachel A. Ridgway, Colin Nixon, Kathryn Gilroy, David M. Gay, Amy Burton, Bin Yan, Katherine Sellers, Vincen Wu, Yuchen Xiang, Engy Shokry, William Clark, Vivian S.W. Li, Simon T. Barry, Richard J.A. Goodwin, Zoltan Takats, Oliver D.K. Maddocks, David Sumpton, Mariia O. Yuneva, Andrew D. Campbell, Josephine Bunch, Owen J. Sansom

## Abstract

With colorectal cancer (CRC) being the second most common cause of cancer-related deaths worldwide^1^, there is an urgent need for better diagnostic tools and new, more targeted therapies. Here we used genetically engineered mouse models (GEMMs), and multimodal mass spectrometry-based metabolomics to study the impact of common genetic drivers of CRC on the metabolic landscape of the intestine. We show that unsupervised metabolic profiling can stratify intestinal tissues according to underlying genetic alterations, and use mass spectrometry imaging (MSI) to identify tumour, stromal and normal adjacent tissues. By identifying ions that drive variation between normal and transformed tissues, we found dysregulation of the methionine cycle to be a hallmark of APC-mutant CRC, and propose one of its enzymes, i.e. adenosylhomocysteinase (AHCY), as a new therapeutic target. Collectively, we show that the profound genotype-dependent alterations in both lipid and small molecule metabolism in CRC may be exploited for tissue classification with no need for ion identification, and we applied further data analysis to expose a novel metabolic vulnerability of CRC.

---

Inactivation of the *APC* tumour suppressor is the most common event in CRC (80%), with co-occurring activation of oncogenic *KRAS (40-50%)*, and/or mutations in other tumour suppressor genes (e.g. *PTEN or TP53*) or oncogenes (e.g. *PIK3CA*) being frequently observed (Figure 1A)^2^. We crossed mice expressing a tamoxifen-inducible intestine-specific Cre recombinase, under the control of the villin promoter (Villin-Cre^ERT2^), with mice harbouring various combinations of conditional alleles of inactivated *Apc* (*Apc^fl^*), *Pten* (*Pten^fl^*), and oncogenic *Kras* (*Kras^G12D^*). Intraperitoneal delivery of tamoxifen resulted in acute gene (in)activation across the intestinal epithelium, with deletion of both copies of *Apc* causing a crypt progenitor phenotype characterised by increased proliferation^3^. To study the metabolic impact of these genetic events, intestinal epithelium was extracted from *wild type* (WT), Villin-Cre^ERT2^ *Kras^G12D/+^* (KRAS), Villin-Cre^ERT2^ *Apc^fl/fl^* (APC), Villin-Cre^ERT2^ *Apc^fl/fl^ Kras^G12D/+^* (APC KRAS), and Villin-Cre^ERT2^ *Apc^fl/fl^ Kras^G12D/+^ Pten^fl/fl^* (APC KRAS PTEN) mice. These were analysed by bipolar forceps Rapid Evaporative Ionizing Mass Spectrometry (REIMS), which allows rapid determination of metabolic profiles with no need for sample preparation and can therefore provide real-time tissue characterization during surgery^4^. The REIMS spectrum is dominated by abundant molecules such as phospholipids, lysolipids, and fatty acids^5, 6^, and accordingly segmentation of the REIMS data by t-distributed stochastic neighbour embedding (t-SNE) was optimal when focusing the analysis on large ions (mass range of *m/z*: 600-1500) (Figure 1B; Figure S1A,B). Inactivation of APC resulted in a clear metabolic differentiation from WT tissues. Additional oncogenic transformation (i.e. APC KRAS and APC KRAS PTEN) resulted in distinct separation (Figure 1B), indicating the profound impact of these clinically relevant events on the metabolic landscape of the intestine. Clustering analysis (*k*-means clustering on t-SNE reduced data, k=5) identified genotype dependent metabolic clusters, and sex-dependent segmentation was observed for samples derived from APC mice as indicated by a male dominant APC cluster (containing 77% of male APC spectra), and female dominant APC cluster (containing 99% of female APC spectra) (Figure S1C). In line with the lack of intestinal hyperproliferation in KRAS mice when sampled at this timepoint^7^, WT and KRAS tissues co-clustered (Figure 1B; Figure S1C). Pathway analysis of the ions discriminating the various genotypes showed that alterations in lipid biosynthesis were largely driving the various metabolic clusters (**Supplemental Tables 1&2)**. Our data reveal a significant effect of APC inactivation and further oncogenic transformation on the intestinal lipidome and highlight the clinical potential of using metabolic phenotyping by REIMS for intrasurgical tissue classification of CRC.

**Figure 1:**
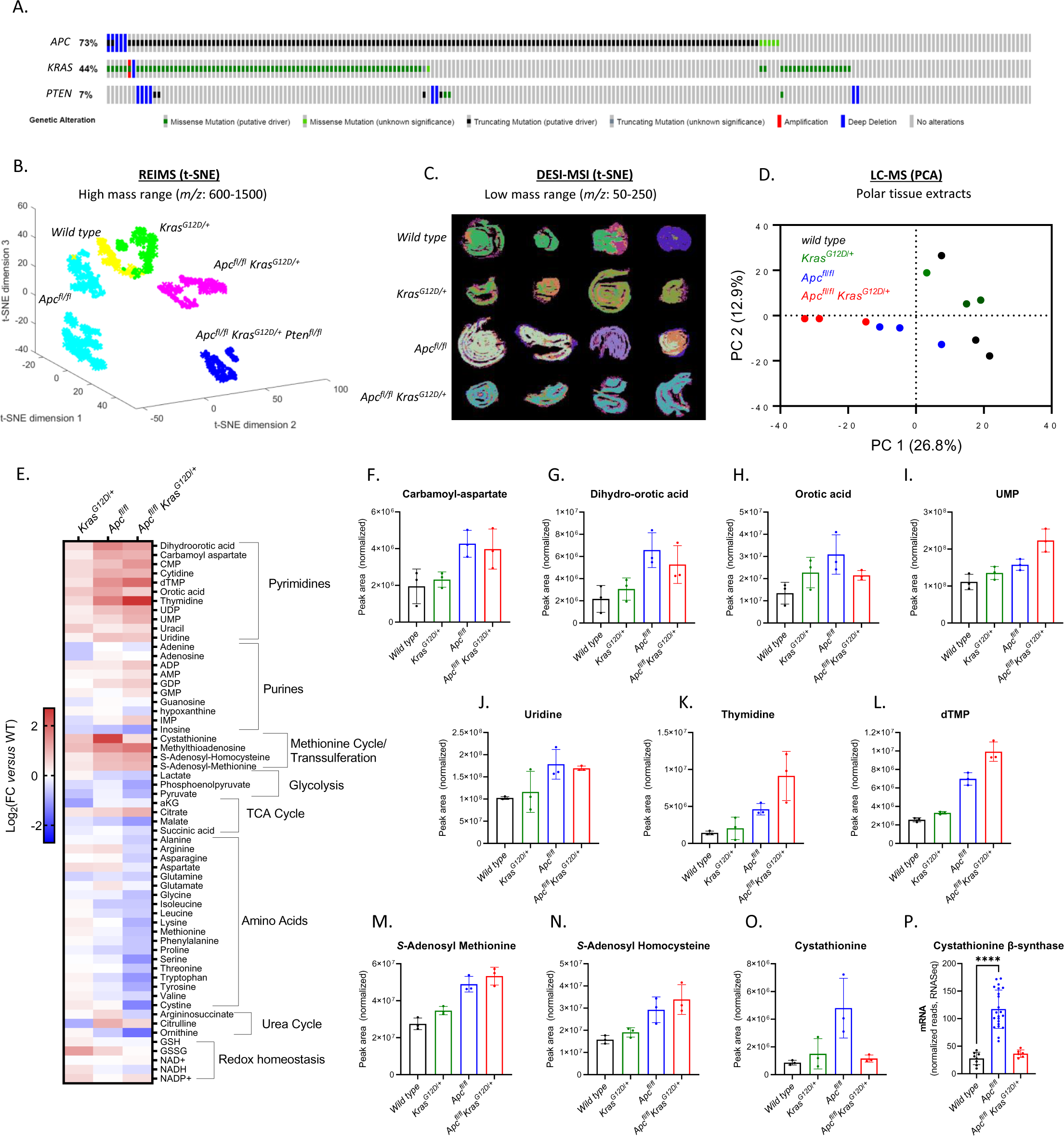
Stratification of GEMMs of intestinal hyperproliferation by metabolic profiling. (A) Oncoprint showing genetic alterations of *APC*, *KRAS*, and *PTEN* in human colorectal adenocarcinoma (TCGA, Firehose Legacy; source: cbioportal.org). (B) t-SNE plot of REIMS data acquired from small intestinal epithelium after specific activation of oncogenic drivers focusing analysis on ions within mass range *m/z*: 600-1500. Each dot corresponds to a single mass spectrum acquired using the REIMS forceps. Data acquired from WT (n=4), KRAS (n=4) APC (n=11), APC KRAS (n=4), and APC KRAS PTEN (n=5) mice. (C) t-SNE of DESI-MSI data acquired from longitudinal sections of the rolled up small intestine of WT, KRAS, APC, and APC KRAS mice focusing analysis on ions within mass range *m/z*: 50-250. (n=4 per genotype, each image represents an individual mouse). (D) PCA of untargeted LC-MS data acquired from polar extracts of small intestinal tissues from WT, KRAS, APC, and APC KRAS mice (n=3 for each genotype). (E) Heatmap showing differences in metabolite abundance in small intestinal tissues of KRAS, APC, and APC KRAS compared with WT mice [targeted analysis of LC-MS data; heatmap constructed based on fold change between the averages of each experimental group (n=3)]. (F-O) Plots showing normalized abundances of (F-H) intermediates of *de novo* pyrimidine synthesis, (I-L) pyrimidine nucleo(s)(t)ides, and (M-O) intermediates of the methionine cycle/transsulferation pathway. (P) Cystathionine β-synthase gene expression as analysed by RNA Sequencing. Asterisk refers to p-value obtained from 2-tailed Mann-Whitney test (****: p<0.0001). Panels F-P show Mean ± SD, each dot represents data from an individual mouse.

REIMS provides a suitable platform for rapid metabolic profiling based on abundant ions in an intact biological specimen but has predominately been employed for lipidomics-based analyses^8, 9^. Electrosurgical REIMS data is largely untested for detection of low mass metabolites with relevance to cancer (e.g. central carbon metabolism, amino acids, and nucleotides) and previously, poor accuracy was found when using the lower end of the mass range due to detection of a large number of fragment ions rather than intact molecules^10^. Small intestinal tissues of WT, KRAS, APC, and APC KRAS mice were therefore additionally analysed using Desorption Electrospray Ionization (DESI) MSI which can provide rich information relating to the spatial distribution of low *m/z* metabolites. As with REIMS, DESI provides an unbiased metabolic readout of intact tissues with no need for solventbased metabolite extraction, which often selectively enriches specific classes of molecules. Similarly, to what is observed with REIMS, untargeted multivariate data analysis focusing the mass range on large ions (*m/z*: 700-1200) showed distinct clustering of APC and APC KRAS, and co-clustering of WT and KRAS tissues (Figure S1D, n_components = 8). However, genotype-dependent clustering was also observed when restricting the analysis to smaller ions (*m/z*: 50-250), indicating additional conserved alterations in non-lipid driven processes which could be used for tissue classification or target identification (Figure 1C, n_components=8). To investigate this further, we analysed the polar fraction of small intestinal tissue extracts by Liquid Chromatography-Mass Spectrometry (LC-MS). Unsupervised PCA analysis showed separate clustering of APC and APC KRAS tissues, and co-clustering of WT and KRAS tissues (Figure 1D). Targeted data analysis revealed increased intermediates of *de novo* pyrimidine synthesis and pyrimidine nucleo(s)(t)ides upon loss of *Apc* (Figure 1E,F-L). Furthermore, we found evidence of increased activity of the methionine cycle as indicated by increased levels of *S*-adenosyl-methionine (SAM), *S*-adenosyl-homocysteine (SAH), and methylthioadenosine (Figure 1E,M,N). Similar regulation of these metabolites was also observed in colonic tissues (Figure S1E). These findings are in line with the previously reported upregulation of pyrimidine synthesis genes by MYC, and increased abundance of SAM in human CRC compared with paired normal tissue^11^. In addition, cystathionine levels (Figure 1O) and gene expression of *Cystathionine ϐ-synthase* (*Cbs*) (Figure 1P) were increased in APC tissues, indicating carbon contribution of homocysteine to the transsulferation pathway. Previous work has shown an association between *KRAS* mutations and epigenetic silencing of *CBS* in primary CRCs^12^, and accordingly we found *Cbs* expression and cystathionine levels decreased in APC KRAS tissues (Figure 1O,P).

These data, obtained in a short-term model of intestinal hyperproliferation, indicate specific metabolic rewiring upon oncogenic transformation of the entire intestine. To test the relevance of our findings in a tumour model, we performed endoscopy-guided injection of 4-OH-tamoxifen in the colonic submucosa of APC and APC KRAS mice (Figure 2A). This resulted in the formation of localized colonic tumours with stromal infiltration (Figure 2B). After confirming tumour development by colonoscopy, the distal colon was dissected and analysed by both DESI-MSI and Matrix-assisted laser desorption/ionization Mass Spectrometry Imaging (MALDI-MSI) to increase metabolite coverage. Multivariate analysis segmented the different tumour compartments (Figure 2C,D; n_components = 15 and 16, respectively), showing that MSI-based metabolic profiling can be applied to detect and classify tumour, stromal and normal tissue in CRC. Certain ions showed pronounced specificity for normal *versus* transformed tissue (Figures S2A, and S3A). Tentative metabolite assignments were validated using a combinatorial approach of tandem mass spectrometry and *in vivo* stable isotope tracing. This revealed specific depletion of glucose (Figure S2) in tumour epithelium compared with stromal or normal tissue. Conversely, we observed specific accumulation of metabolites in tumour epithelium (e.g. *m/z* 464.288 for which MSMS data suggest lysophophatidylethanolamine (17:1) as a likely assignment; Figure S3).

**Figure 2:**
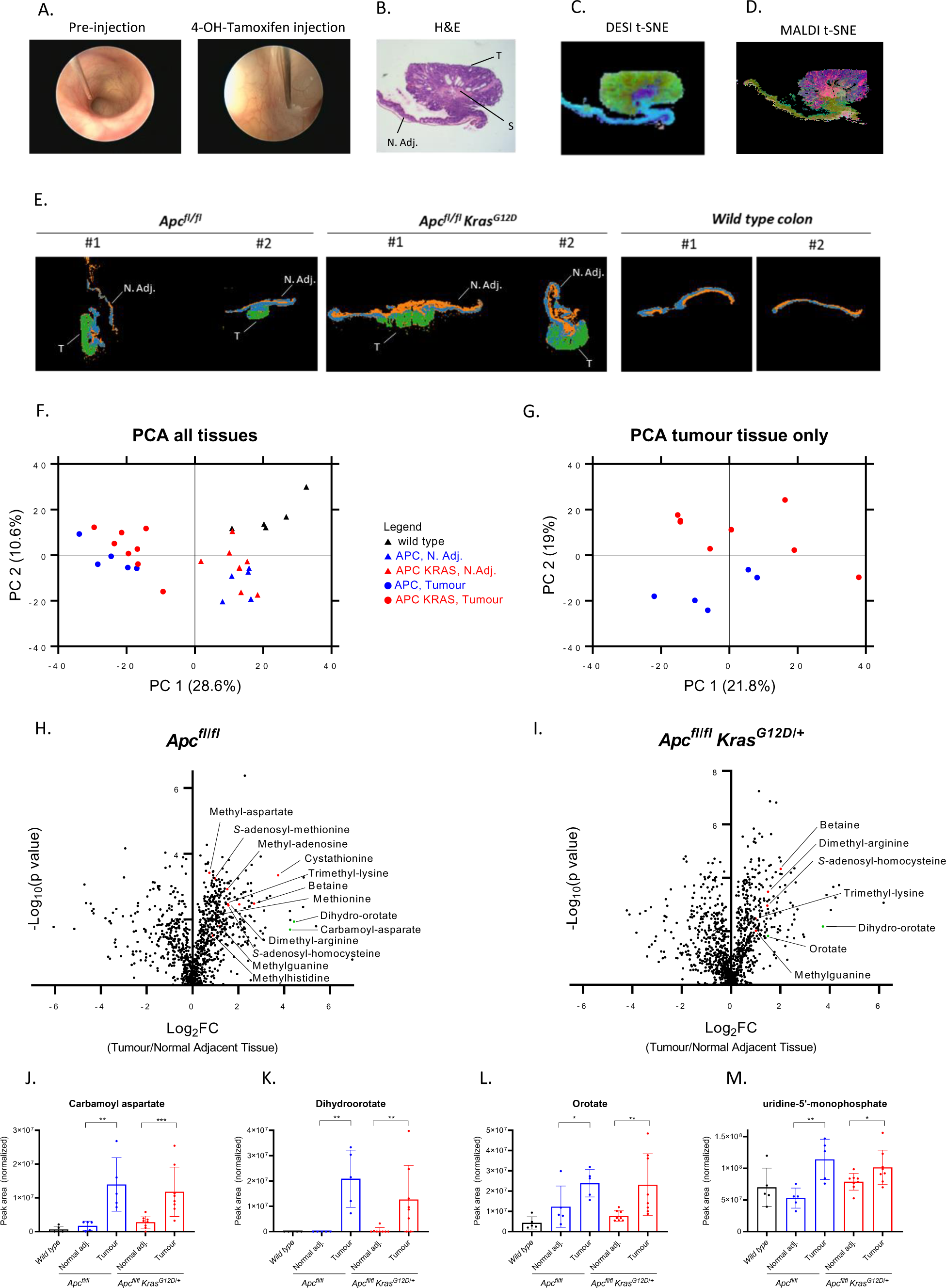
Untargeted metabolic profiling of GEMMs of APC-deficient CRC using MSI and LC-MS. (A) Representative images of endoscopy-guided submucosal delivery of 4-OH-tamoxifen in the murine colon resulting in localized genetic recombination and tumour formation. (B) H&E of distal colon tissue with an Apc-deficient tumour [N. Adj.: normal adjacent tissue; S: stroma; T: tumour tissue]. (C) tSNE plot of data acquired by DESI-MSI (negative polarity) and (D) MALDI-MSI (negative polarity) of distal colon tissue of locally induced APC mice (tissues analysed for n=4 animals; n=1 shown). (E) tSNE plots of data acquired by DESI-MSI (positive polarity) of distal colon tissue of locally induced APC (n=5) and APC KRAS (n=7) mice, and normal colon from WT (n=4) mice (n=2 mice per group shown). (F) PCA of untargeted LC-MS data acquired on polar extracts of normal adjacent and colon tumour tissues of APC (n=5) and APC KRAS (n=8), and control colon of WT (n=5) mice. (G) PCA of untargeted LC-MS data acquired on polar extracts of tumour tissues of APC (n=5) and APC KRAS (n=8) mice. (H,I) Volcano plots showing metabolic differences between paired normal adjacent and colon tumour tissues of APC (n=5) or APC KRAS (n=8) mice as detected by untargeted LC-MS (p-values obtained from paired t-tests. All annotated metabolites: FC≥1.5 and significant after Benjamini-Hochberg FDR correction (q=0.05); Red dots: metabolites related to methionine metabolism; Green dots: intermediates of *de novo* pyrimidine biosynthesis. (J-M) Plots showing normalized abundances (Mean ± SD) of intermediates of *de novo* pyrimidine synthesis in tumour and normal adjacent colon tissue of APC (n=5 mice) and APC KRAS (n=8 mice), and control colon of WT (n=5) mice (targeted analysis of LC-MS data). Each dot represents an individual mouse, asterisks refer to p-values obtained from 1-tailed Mann-Whitney tests (*: p<0.05; **: p<0.01; ***: p<0.001).

Multivariate analysis of data acquired on large colonic sections containing both tumour and normal adjacent tissue (NAT) did not distinguish between APC and APC KRAS tumours (Figure 2E), indicating that the observed metabolic differences between tumour and NAT exceed the impact of oncogenic *Kras* expression. This was confirmed by untargeted LC-MS analysis of colon tumour tissue and paired NAT from tumour-bearing animals, and control colonic tissues from WT mice (Figure 2F). However, restricting the PCA to tumour tissues only resulted in genotype-dependent segmentation, stratifying tumours according to *Kras* status (Figure 2G). This indicates that careful spatial analysis of the various tumour compartments is key to understanding the metabolic impact of oncogenic drivers such as *Kras*. Therefore, we compared APC KRAS *versus* APC colon tumours using both DESI-MSI and LC-MS. We focused analysis of the data acquired by DESI-MSI on tumour epithelial regions only, whereas LC-MS was performed on bulk tumour tissue. Both modalities revealed specific metabolic effects of KRAS^G12D^ (Figure S4A,C; Supplemental Tables 3&4). DESI-MSI exposed a specific decrease in glutamine in APC KRAS tumour epithelium (Figure S4A,B), which was not revealed by untargeted LC-MS (Figure S4C). We recently reported the amino acid transporter SLC7A5, importing large neutral amino acids at the expense of intracellular glutamine, to be a targetable vulnerability of KRAS-mutant CRC^7^. Our current data exemplify the value of spatial metabolomics using MSI to complement bulk analysis of tumour tissue where metabolites from different compartments (i.e. tumour epithelial, stromal, immune, and normal adjacent cells) are mixed.

We next focused on tumour-specific polar metabolic alterations by analysing paired tumour and normal adjacent colonic tissues. This revealed increased levels of dihydroorotate (Figure 2H,I), other intermediates of *de novo* pyrimidine synthesis, and pyrimidine nucleotides in APC and APC KRAS tumours (Figure 2J-M). Also, there was a striking tumour-specific enrichment in methionine-related metabolites, and methylated metabolites requiring SAM (Figure 2H,I). These data confirmed what we observed in the acute model of intestinal hyperproliferation (Figure 1E-O). Given the well-described effects of interfering with pyrimidine biosynthesis and salvage in CRC (e.g. by antimetabolites), we focused our attention on the methionine cycle (Figure 3A). Targeted data analysis showed that SAM and SAH levels were significantly increased in tumour compared to paired NAT of both APC and APC KRAS mice (Figure 3B,C). Also, there was a tumour-specific increase in cystathionine with lower cystathionine levels being observed in APC KRAS compared to APC tumours (Figure 3D). We analysed publicly available transcriptomic data from TCGA (Pancancer Atlas) to understand which enzymes of the methionine cycle are transcriptionally regulated in human CRC, and found increased adenosylhomocysteinase (*AHCY*; or SAH hydrolase) expression in colorectal adenocarcinoma compared to normal colon tissue (Figure 3E). ACHY converts SAH into homocysteine, and its expression is regulated by MYC^13^. AHCY activity is required for MYC-induced mRNA Cap methylation^13^, and its gene expression correlates with DNA methylation state in tumours^14^. Pan-cancer analysis across 17 different cancer types showed highest expression of *AHCY* in CRC (Figure S5A). Molecular profiling of human CRC revealed enrichment of *ACHY* expression in consensus molecular subtype 2 (CMS2) (Figure 3F), which is characterized by WNT and MYC activation, and accounts for 37% of CRCs^15^. We found acute loss of *Apc* to be sufficient to drive *Ahcy* overexpression in the murine small intestine (Figure 3G). AHCY protein expression was found enriched in intestinal crypts of WT animals as shown by immunofluorescence (IF) (Figure 3H). Furthermore, high AHCY expression was also observed in adenomas of *Apc^Min/+^* mice (Figure 3H), and in *Apc*-deficient compared with WT organoids (Figure 3I). Analysis of publicly available single-cell RNA sequencing data of the murine small intestinal epithelium^16^ confirmed *Ahcy* expression to be highest in early enterocyte progenitor cells (Figure 3J). Notably, IF also showed increased levels of 5-methylcytosine in *Apc*-deficient organoids, which requires SAM as a methyl donor and therefore methionine cycle activity (Figure S5B).

**Figure 3.**
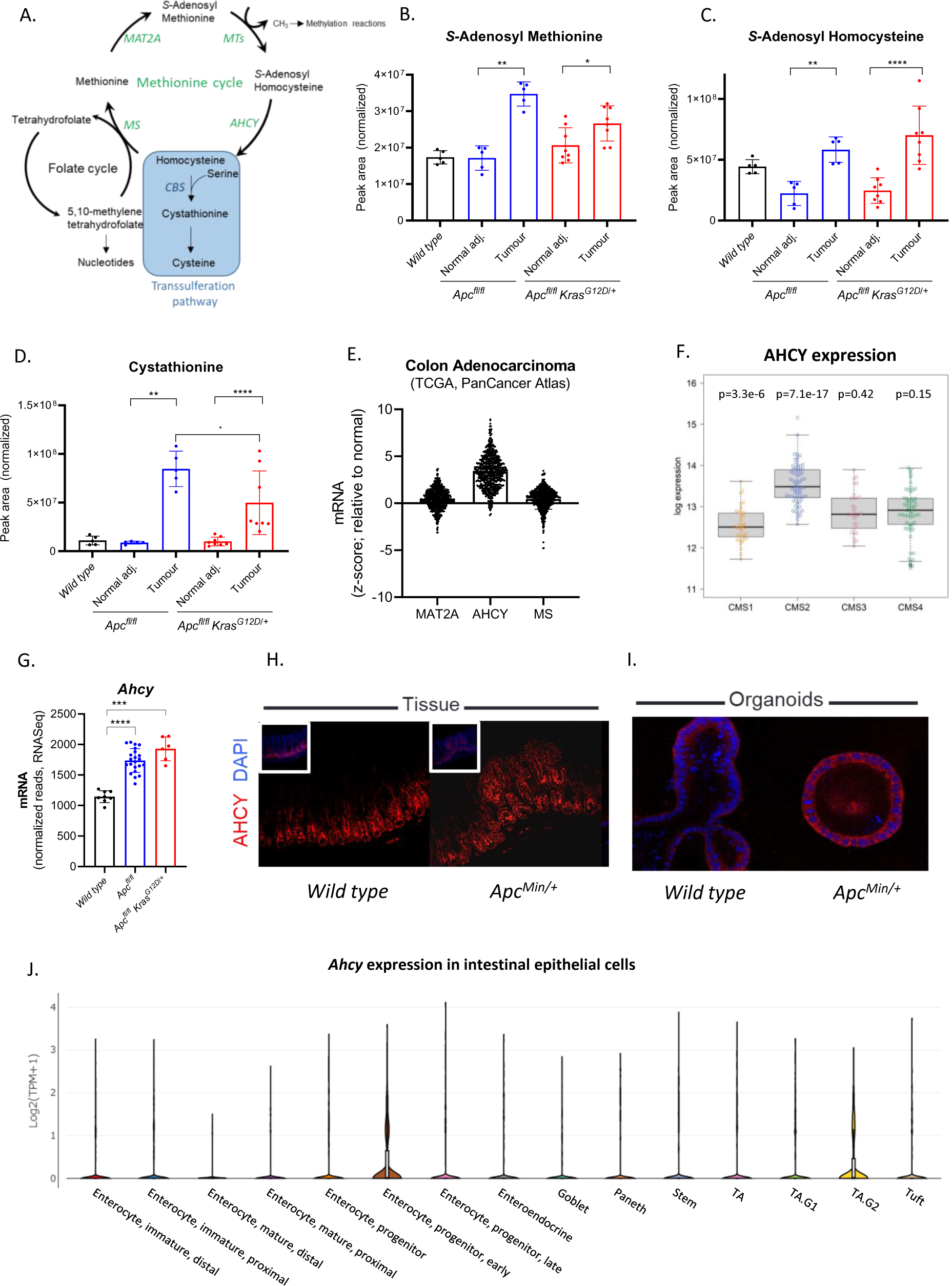
Methionine cycle activity and AHCY expression is increased in CRC. (A) Schematic representation of the methionine and folate cycle, and transsulferation pathway. (B-D) Plots showing normalized abundances (Mean ± SD) of (B) SAM, (C) SAH, and (D) cystathionine in tumour and normal adjacent colon tissue of APC (n=5) and APC KRAS (n=8), and control colon of WT (n=5) mice as detected by targeted analysis of LC-MS data. Each dot represents an individual mouse, asterisks refer to p-values obtained from 1-tailed Mann-Whitney tests (*: p<0.05; **: p<0.01; ****: p<0.0001). (E) Expression of genes encoding enzymes of the methionine cycle in human colorectal adenocarcinoma compared with normal colon (TCGA, PanCancer Atlas; source; cbioportal.org). (F) Expression of *AHCY* across the different consensus molecular subtypes of human CRC^15^. p values refer to difference between each CMS group relative to all others. (G) *Ahcy* expression (Mean ± SD) in the small intestine of WT (n=8), APC (n=22) and APC KRAS (n=6) mice. Each dot represents an individual mouse, asterisks represent p-values obtained from 1-tailed Mann-Whitney tests (***: p<0.001; ****: p<0.0001). (H-I) IF showing AHCY protein expression in the small intestine and organoids derived from the intestine of WT and adenomas from *APC^Min/+^* mice. (J) *Ahcy* gene expression across the different cell population of the murine small intestine^16^ (source: https://singlecell.broadinstitute.org/).

Dietary restriction of methyl donors has been shown to reduce tumour burden in *Apc^Min/+^ mice*^17^. The methionine cycle also supports the regeneration of tetrahydrofolate via methionine synthase (MS) and recently, MS was found essential to support nucleotide synthesis and tumour cell proliferation under physiological levels of folate^18, 19^. In addition, methionine restriction sensitizes to chemotherapy and radiation by disrupting one-carbon metabolism^20^. Our data indicate functional activity of the methionine cycle in APC-mutant CRC, and that AHCY may be important for its regulation. To test the effect of pharmacological AHCY inhibition, crypt cells were isolated from the small intestine of APC mice and cultured *in vitro* as organoids. Treatment with the AHCY inhibitor 3-deazaneplanocin A (DZNeP)^21–23^ significantly impaired organoid growth (Figure 4A,B). To understand the metabolic effects of AHCY inhibition in these organoids, we traced the fate of ^13^C_5_-methionine in the presence/absence of DZNeP (Figure 4C). Whereas the intracellular levels of methionine or SAM were not affected (Figure 4D,E; Figure S6A,B), we observed a pronounced increase in SAH, the substrate of AHCY, upon treatment (Figure 4F; Figure S6C). DZNeP decreased labelling of trimethyllysine from methionine (Figure 4G; Figure S6D). This demonstrates that AHCY inhibition reduces methyltransferase activity which may affect DNA/RNA, protein and metabolite methylation. We could not detect homocysteine in these samples but found the intracellular levels of cystathionine to be markedly reduced upon treatment with DZNeP (Figure 4H; Figure S6E) indicating a decreased contribution of methionine-derived carbons to the transsulferation pathway.

**Figure 4.**
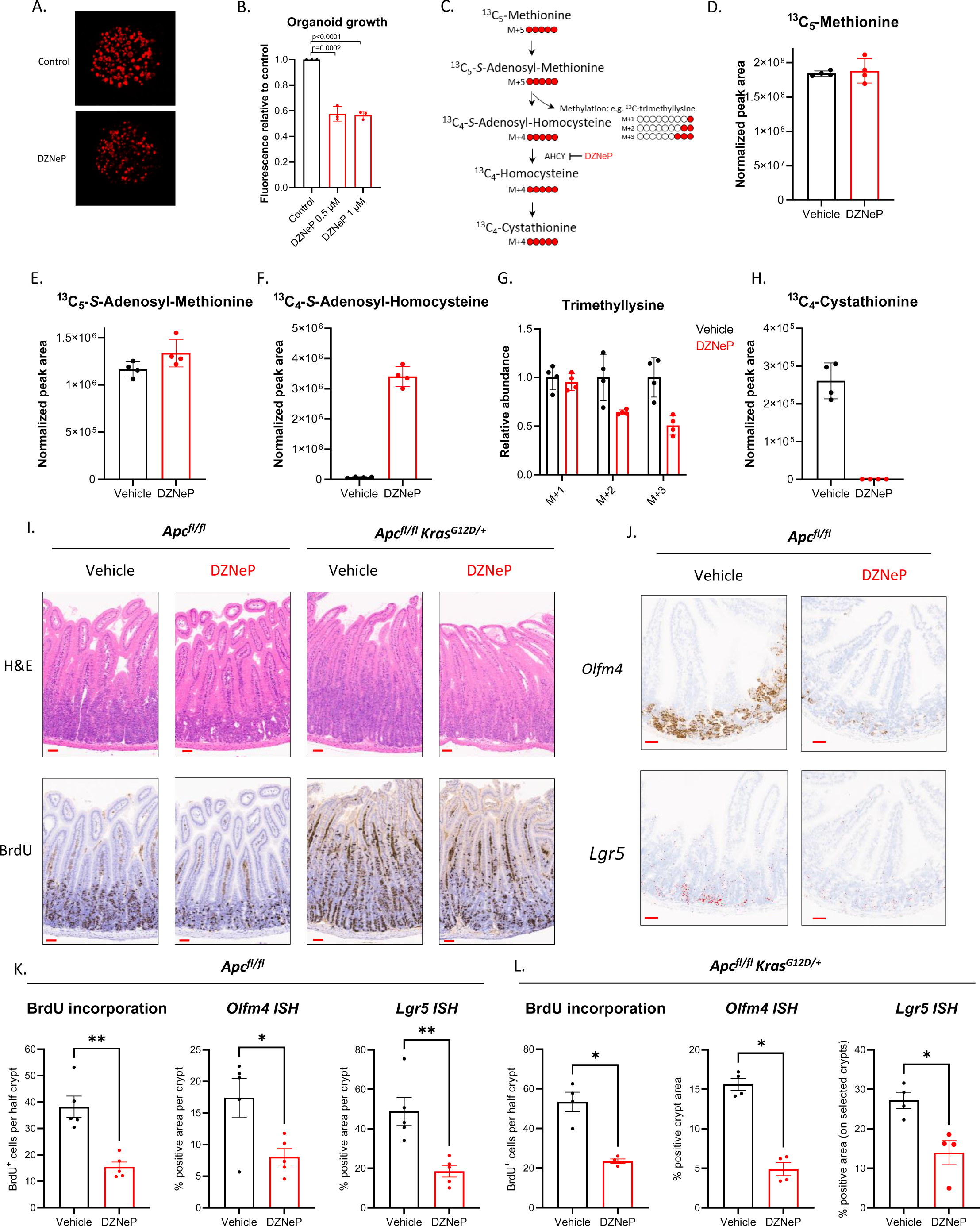
Inhibition of AHCY suppresses proliferation and stem cell expansion in APC and APC KRAS driven models of CRC. (A) Representative image and (B) quantification of APC organoids (+/- DZNeP 1 μM) stained with Syto 60 Nucleic Acid Stain (Mean ± SD; each dot represents the mean of 3 independent experiments with 4 technical replicates each; p values generated using an unpaired, two-tailed t-test). (C) Schematic showing carbon contribution of ^13^C_5_-methionine to intermediates of the methionine cycle, cystathionine and trimethyllysine. Abundance of (D) ^13^C_5_-methionine, (E) ^13^C_5_-SAM, (F) ^13^C_4_-SAH, (G) various isotopologues of trimethyllysine, and (H) ^13^C_4_-cystathionine in APC organoids (+/- DZNeP 1 μM) (Mean ± SD; data from a representative experiment performed twice, with 4 technical replicates each; each dot represents a technical replicate). (I) Representative images of H&E staining and IHC for BrdU on small intestinal sections of APC (n=5) and APC KRAS (n=4) mice treated with vehicle or DZNeP (5 mg/kg). Scale bars: 50 μm; (J) Representative images of ISH for *Olfm4* and *Lgr5* expression in the small intestine of APC mice (n=5) treated with vehicle or DZNeP (5 mg/kg). Scale bars: 50 μm; (K,L) Quantification of IHC for BrdU and ISH for *Olfm4* and *Lgr5* in the small intestine of APC and APC KRAS mice treated with vehicle or DZNeP (5 mg/kg). [Mean ± SEM; n=5 (APC) or n=4 (APC KRAS) mice per experimental arm; BrdU: each dot represents the average number of BrdU positive cells per half crypt for each mouse; *Olfm4* and *Lgr5*: each dot represents the average % positive area per crypt for each mouse. Asterisks refer to p-values obtained from 1-tailed Mann-Whitney tests (*: p<0.05; **: p<0.01)].

To study the effect of AHCY inhibition *in vivo*, APC and APC KRAS mice were treated with DZNeP post intraperitoneal induction with tamoxifen (Figure S7A&B). The APC protein is a component of the β-catenin destruction complex and plays a critical role in maintenance of the stem cell niche and tumour suppression in the intestinal epithelium. APC negatively regulates WNT/TCF signalling by directing the degradation of β-catenin^24^. *APC* deficiency results in accumulation of nuclear β-catenin and thereby increasing transcription of its target genes (including *c-MYC* and *AXIN2)*^25, 26^. Previously, we have shown that acute loss of *Apc* leads to a crypt progenitor phenotype with increased proliferation, stem cell markers, and perturbed differentiation^27, 28^. DZNeP did not affect direct WNT pathway activation as analysed by immunohistochemical analysis of nuclear accumulation of β-catenin in APC mice (Figure S7C) but significantly suppressed intestinal hyperproliferation in both the small intestine (Figure 4I,K,L; Figure S7D,E) and colon (Figure S7F-J), without impairing intestinal crypt proliferation in *wild type* animals (Figure S8A-D). Given the high expression of AHCY in intestinal crypts, we analysed the expression of intestinal stem cell markers using *in situ* hybridization in APC and APC KRAS mice. Expression of both *Olfm4* and *Lgr5* was significantly reduced upon DZNeP treatment (Figure 4J,K,L; Figure S8E), indicating that AHCY inhibition prevents the crypt progenitor phenotype driven by *Apc* loss, both in the presence and absence of *Kras^G12D^* expression. Collectively, these data show that AHCY inhibition truncates the methionine cycle and thereby reduces proliferation of APC-deficient cells indicating its potential as a new actionable target for APC-driven CRC, even in the context of mutant KRAS.

In conclusion, we applied multi-modal mass spectrometry-based metabolomics to investigate the metabolic consequences of common oncogenic events in CRC. We show that various combinations of genetic alterations in *Apc*, *Kras*, and *Pten* perturb intestinal epithelial metabolism to the extent that metabolic profiling can accurately stratify tissues according to underlying genetic events, with no need for further ion identification. We used REIMS as a tool for rapid lipidomic segmentation highlighting the clinical potential of our results, and applied MSI to classify tumour epithelial, tumour stromal, and normal adjacent tissues. Finally, untargeted LC-MS indicated increased activity of the methionine cycle in APC-mutant CRC, which ultimately revealed AHCY as a promising target for CRC. Our results argue for using a combination of these various analytical platforms to study metabolic rewiring in cancer.

## Methods

### Mouse studies

All *in vivo* experiments were carried out in accordance with the UK Home Office regulations (under project licences: 70/8646 and PP3908577 and P609116C5), and by adhering to the ARRIVE guidelines with approval from the Animal Welfare and Ethical Review Board of the University of Glasgow and the Francis Crick Institute. Mice were housed under a 12 h light-dark cycle, at constant temperature (19-23 °C) and humidity (55 ± 10%). Standard diet and water were available *ad libitum*. The majority of the work was performed in the C57BL/6J background. The following alleles were used in this study: *VillinCreER*^29^, *Apc^fl^*^30^, *Kras^G12D^* ^31^, *Pten^fl^*32 and *Apc^Min^* ^33^. For conditional alleles, robust recombination throughout the intestinal epithelium was obtained by one or two i.p. injections of 2 mg tamoxifen, and tissues were harvested 3 or 4 days post induction. To drive spatially localized Cre induction in the colon, 4-hydroxytamoxifen (70 μL, 100 nM; Merck Millipore) was delivered under general anaesthesia via a single injection into the colonic submucosa via colonoscopy as described by Roper *et al*.^34^. Tumour formation was confirmed via colonoscope prior to tissue sampling. Animals heterozygous for *Apc^Min^* allele of both sexes at the age of 12-16 weeks were used. Samples from *Apc^Min/+^* mice were collected at the onset of clinical signs of intestinal adenomas. For tissue metabolomics (MSI or LC-MS), intestines were flushed with ice-cold PBS, tissues of interest were dissected and snap frozen.

To study the effect of AHCY inhibition *in vivo,* Villin-Cre^ERT2^ *Apc^fl/fl^* (APC) and Villin-Cre^ERT2^ *Apc^fl/fl^ Kras^G12D/+^* (APC KRAS) mice were treated with DZNeP.HCl (5 mg/kg i.p.; Carbosynth) or vehicle (PBS) from day 1 post i.p. tamoxifen administration (Figure S7A,B). Animals were injected with BrdU (i.p.) 2 hours prior to sampling tissues.

To study tumour-specific distribution of glycocholic acid (GA; **Figure S3**), ^13^C-GA (CLM-191, CK Isotopes) was administered by oral gavage to tumour-bearing animals (APC, spatially localized Cre induction) at 75 mg/kg (vehicle: 10% DMSO, 0.5% hydroxypropyl methylcellulose (HPMC) + 0.1% Tween-80). Tissues were harvested and snap frozen for downstream analysis 8.5 hours post administration.

### Haematoxylin & Eosin, Immunohistochemistry and RNA-ISH

Intestinal tissues were fixed in 10% neutral buffered formalin or methacarn (methanol, chloroform, and acetic acid; 4:2:1 ratio). All haematoxylin & eosin (H&E), immunohistochemistry (IHC) and *in situ* hybridisation (ISH) staining was performed on 4µm formalin fixed paraffin embedded sections (FFPE) which had previously been heated at 60⁰C for 2 hours.

Standard protocols were used for H&E staining. FFPE sections for BrdU (347580, Becton Dickinson) IHC staining were loaded into an Agilent pre-treatment module to be dewaxed and undergo heat induced epitope retrieval (HIER) using high pH target retrieval solution (TRS) (K8004, Agilent). The sections were heated to 97⁰C for 20 minutes in high pH TRIS buffer. After HIER the sections were rinsed in flex wash buffer (K8007, Agilent) prior to being loaded onto the Agilent autostainer. Mouse on mouse blocking reagent (MKB-2213, Vector Labs) was applied to the sections for 20 mins before washing with wash buffer. The sections underwent peroxidase blocking (S2023, Agilent) for 5 minutes and were then rinsed with flex buffer before applying BrdU antibody to the sections at a previously optimised dilution (1/250) for 35 minutes. The sections were washed with flex wash buffer before application of mouse envision secondary antibody (K4001, Agilent) for 30 minutes. Sections were rinsed with flex wash buffer before applying Liquid DAB (K3468, Agilent) for 10 minutes. The sections were washed in water and counterstained with haematoxylin z (RBA-4201-00A, CellPath).

ISH for *Lgr5* (312178) and *Olfm4* (311838) (both from Advanced Cell Diagnostics) was performed using RNAscope 2.5 LSx (brown) detection kit (322700, Advanced Cell Diagnostics) on a Leica BOND Rx autostainer strictly according to the manufacturer’s instructions. Images were analysed using HALO software (Indica Labs).

### Immunofluorescence

Intact intestinal tissues were flushed with PBS followed by 10% buffered formalin (Sigma-Aldrich). Intestines were then fixed as a swiss roll in 10% buffered formalin over night with gentle rocking. The tissues were rinsed in 70% ethanol before embedding in paraffin wax. 4 µm sections were dewaxed and rehydrated by 2-time immersion in xylene for 10 minutes, then 2-time immersion in 100% ethanol for 10 minutes, followed by subsequent immersion in 95, 70, 50, and 25% ethanol for 5 minutes, and rinsed in water. Antigen retrieval was performed for 20 minutes at high temperature with 10 mM citrate, pH 6.0. Slides were washed with PBS and blocked with 1% bovine serum albumin (BSA), 0.1% triton X-100 in PBS.

Organoids were seeded in 8-well slide chambers (Thermo Scientific Nunc 154526) and fixed in prewarmed 4% formalin for 25 min and then rinsed and permeabilized by incubating for 1 hour in 1% BSA, 0.1% triton X-100 in PBS.

Tissue or cell culture slides were incubated with an appropriate dilution of primary antibody [rabbit anti-AHCY (ProteinTech 10757-2-AP; 1:100) and mouse anti-5m-Cystosine (Abcam Ab10805; 1:100) in a humidity chamber overnight at 4°C. Slides were washed 5 times in PBS for 5 minutes with gentle rocking, followed by a 2 hour incubation with 1:250 dilution of either anti-rabbit conjugated to alexa 555 or anti-mouse conjugated to alexa 488, as appropriate, in blocking solution for 1 hour, and washed again 5 times in PBS for 5 minutes with gentle rocking. Slides were mounted to a coverslip with Vectashield (Vector Laboratories) containing 0.1 µg/ml DAPI. Images were captured with a Leica SP5 confocal microscope.

### RNA Sequencing

For RNA sequencing 1 µg of RNA was prepared in 50 µl. RNA sequencing was performed using an Illumina TruSeq RNA sample prep kit, then run on an Illumina NextSeq using the High Output 75 cycles kit (2 x 36 cycles, paired end reads, single index). The raw sequence quality was assessed using the FastQC algorithm version 0.11.8. Sequences were trimmed to remove adaptor sequences and low-quality base calls, defined by a Phred score of <20, using the Trim Galore tool version 0.6.4. The trimmed sequences were aligned to the mouse genome build GRCm38.98 using HISAT2 version 2.1.0, then raw counts per gene were determined using FeatureCounts version 1.6.4. Differential expression analysis was performed using the R package DESeq2 version 1.22.2, and principal component analysis was performed using R base functions.

### Organoid studies

Intestines from wild type C57Bl/6J mice were sliced longitudinally, rinsed in PBS-CMF (calcium and magnesium free PBS) with vigorous shaking, and cut into 2 cm segments. Villi were scraped and rinsed away. The tissue was minced and collected in a falcon tube. The tissues were washed approximately 10 times by resuspension in fresh PBS-CMF, pelleting by gravity, aspiration and resuspension in fresh PBS-CMF. The tissue was centrifuged at 1000 rpm for 5 minutes. PBS-CMF was aspirated. Crypts were removed by suspension in 10 mL 2 mM EDTA and gentle agitation for 30 minutes followed addition of 10 mL PBS-CMF and vigorous shaking. EDTA was neutralized by 5 mL PBS with calcium and magnesium. The Crypt-free tissue was allowed to settle to the bottom of the tube and the crypt-containing supernatant was collected. Crypts were pelleted and washed 2 x with Advanced DMEM-F12 by centrifugation at 1000 rpm for 5 minutes. The crypts were resuspended in matrigel and seeded as drops onto culture dishes. After the matrigel solidified, 1:1 Advanced DMEM-F12 media with 10 mM HEPES, 1x B27, 1 mM N-acetylcysteine, 50 ng/mL EGF, 500 ng/mL R-spondin, and 200 ng/mL Noggin: WNT conditional media^35^ was added. After the crypts formed spheres, the WNT conditional media was removed, which allowed the spheres to differentiate and bud.

APC^Min^ adenomas were isolated by slicing the adenoma-bearing tissues longitudinally, rinsing with PBS-CMF, and excising adenomas with forceps. Adenomas were dissociated by incubating with 1 mg/mL collagen dispase in Advanced DMEM-F12 for 1 hour. Dispase was neutralized with 5% FBS. Single cells were isolated by filtering and pelleting by centrifugation at 1000 rpm for 5 minutes. The cells were resuspended in matrigel and seeded as drops onto culture dishes. APC-mutant cells were selected by not including WNT or R-spondin in the media. After the establishment of organoids, organoids were cultured in Advanced DMEM-F12 media with 10 mM HEPES, 1x B27, 1 mM N-acetylcysteine, 50 ng/mL EGF, 500 ng/mL R-spondin, and 200 ng/mL Noggin.

Organoids were isolated from the small intestine of *VilCreER Apc^fl/fl^* mice as described previously^36^. Organoids were resuspended in Matrigel (BD Bioscience), plated in 24 well plates and supplemented with Advanced DMEM/F12 supplemented with 10 mM HEPES, 2 mM glutamine, N2, B27 (all from Gibco, Life Technologies), 100 ng/ml Noggin, and 50 ng/ml (both from PeproTech).

To evaluate the effect of AHCY inhibition on growth, organoids were seeded as fragments in Matrigel and cultured in the presence/absence of DZNeP.HCl (0, 0.5 or 1μM) for 72 hours. Culture medium was removed and organoids were stained for 90 min with SYTO^TM^ 60 Nucleic Acid Stain (0.5 μM; Thermo Fisher). Fluorescent signal was quantified on a LICOR imaging system.

To study the metabolic effects of AHCY inhibition, organoids were seeded as fragments in Matrigel, supplemented with Advanced DMEM/F12 (containing 10 mM HEPES, 2 mM glutamine, N2, B27, 100 ng/ml Noggin, and 50 ng/ml) and allowed to proliferate for 48 hours. Next, organoids were pre-treated for 2 hours with DZNeP.HCl (0 or 1μM final concentration), after which medium was replaced with fresh medium supplemented with ^13^C_5_-methionine (116 μM) and DZNeP.HCl (0 or 1μM final concentration). After 18 hours, cells were washed 3 times with ice cold PBS and extracted in 400 μL extraction solution (methanol:acetonitrile:water; 50:30:20 ratio). Samples were centrifuged and analysed by LC-MS on a ZIC-pHILIC column as described below. Extracted organoids were stained with SYTO^TM^ 60 Nucleic Acid Stain as described above and fluorescent signal was used for data normalization.

### HPLC-MS

Frozen tissue fragments were weighed and homogenized in ice-cold extraction solution (20 mg/mL) using ceramic beads and a Precellys Homogenizer (Bertin Instruments). Samples were centrifuged (10 min, 16,000*g*) and the supernatant was analysed as described below.

A Thermo Ultimate 3000 high performance liquid chromatography (HPLC) system was equipped with a ZIC-pHILIC column (SeQuant; 150 mm by 2.1 mm, 5 μm; Merck KGaA, Darmstadt, Germany), with a ZIC-pHILIC guard column (SeQuant; 20 mm by 2.1 mm) for metabolite separation. Cell or tissue extracts were injected (5 μL) and metabolite separation was obtained as described before^37^. The HPLC system was coupled with a Q Exactive Plus Orbitrap Mass Spectrometer (Thermo Fisher Scientific) used with a resolution of 70,000 at 200 mass/charge ratio (*m/z*), electrospray ionization, and polarity switching mode across a mass range of 75 to 1000 *m/z*. Mass accuracy was below 5ppm. Untargeted metabolomics was performed as previously described^37^. In brief, where required for metabolite identification, a mixture of all samples within an experiment was analysed in both positive and negative single ionisation mode using data-dependent fragmentation (ddMS2). Data was acquired using Xcalibur software (v4.3, Thermo Scientific). Untargeted data analysis was performed using Compound Discoverer (v3.2, Thermo Scientific). Retention times were aligned across all sample data files (maximum shift 2 min, mass tolerance 5 ppm). Unknown compound detection (minimum peak intensity 5e5) and grouping of compound adducts were carried out across all samples (mass tolerance 5 ppm, RT tolerance 0.7 min). Missing values were filled using the software’s Fill Gap feature (mass tolerance 5 ppm, S/N tolerance 1.5). Metabolite identification was achieved by matching the mass and retention time of observed peaks to an in-house database generated using metabolite standards (mass tolerance 5 ppm, RT tolerance 0.5 min), Peak annotations were confirmed using mzCloud (ddMS2) database search (precursor and fragment mass tolerance of 10 ppm, match factor threshold 50) and searching predicted compositions (mass tolerance 5 ppm, minimum spectral fit and pattern coverage of 30% and 90%, respectively) against the HMDB database. Targeted data analysis was performed using Tracefinder (v4.1, Thermo Scientific). Statistical tests were performed using Compound Discoverer, Perseus 1.6.2.2^38^ and Graphpad Prism 9.

#### Detection of ^12^C-, and ^13^C-glycocholic acid

Single reaction monitoring (SRM) mode was used to detect glycocholic acid on a Altis QQQ Mass Spectrometer equipped with a Vanquish LC system (Thermo Fisher Scientific). Separation of metabolites was performed on a Acquity HSS T3 column (Waters; 150 mm by 2.1 mm, 1.8 μm). The mobile phase consisted of solvent A (water with 0.1% formic acid) and solvent B (acetonitrile with 0.1% formic acid) using the following gradient: 0 min 20% B, 8 min 95% B, 10 min 20% B at a constant flow rate of 0.3 mL/min. Injection volume was 5 µL. Two transitions were optimised using an authentic standard of glycocholic acid (glycine-1-^13^C, CLM-191-PK, Cambridge Isotope Laobratories, UK), from the negative precursor ion (*m/z* 465) to product ions (*m/z* 402, and *m/z* 75). Total cycle time was 0.8 seconds and Q1 resolution was (FWHM) 0.7 and Q3 (FWHM) 1.2. For each transition, the collision energy applied was optimized to generate the greatest possible signal intensity. The optimized source parameters were spray voltage 2500V, sheath gas 35, Aux gas 7, ion transfer tube temp 325 C, Vaporizer temp 275 C, RF lens 105. Data acquisition was performed using Xcalibur 4.1 software.

### REIMS

REIMS was performed on intestinal epithelium extracts, prepared as previously described^39^. An Erbe VIO 50C electrosurgical generator (Erbe Elektonedizin, Tübingen, Germany) operated in bipolar mode at 25 W was used to power the sampling forceps. Samples were allowed to reach room temperature prior to sampling, a portion of each pellet was removed using one tip of the forceps, the two electrodes of the forceps were brought into close proximity and the forceps were activated using a foot switch. The generated smoke was aspirated and directed to the REIMS interface using a 2 m long Tygon tube. A Waters isocratic solvent manager (Waters, UK) was used to introduce propan-2-ol to the Venturi of the REIMS interface at a flow rate of 0.1 mL/min, where it was mixed with the aspirated aerosol. REIMS data were acquired in negative ion mode (50-1500 *m/z*) using a Waters Xevo G2-XS QToF mass spectrometer fitted with a REIMS interface (30,000 mass resolving power at *m/z* 956, < 1 ppm RMS mass accuracy; Waters Corporation, UK).

Data were converted from proprietary .RAW format to imzML using ProteoWizard^40^ and imzML Converter^41^; and analysed using SpectralAnalysis^42^ in MatLab (2019b). Data were preprocessed using interpolation rebinning with a bin with of 0.001 Da, to create a consistent *m/z* axis to generate a mean spectrum. The mean spectrum was then peak picked using a gradient method, and the top 2000 peaks selected for further processing. The regions where the forceps were active were then identified by clustering the data using k-means clustering using the cosine distance with k=2. Data were then l_2_ normalised prior to further analysis by t-SNE and clustering.

t-SNE was applied to these data using the Matlab function “tsne” (Matlab v2017a, Statistics and Machine Learning Toolbox) reducing the data to 3 dimensions using the cosine distance metric and the default hyperparameters^43^. Clustering was then performed on the t-SNE reduced data using the Matlab “kmeans” function using the Euclidean distance and 10 replicates and 5 clusters (5 clusters were chosen as there were 5 genotypes included in the study).

Following the segmentation, pairwise mean intensity log_2_ fold change and two sided t-test p-values were calculated between the data for each *m/*z from each genotype. The discriminating ions from this analysis (absolute log_2_ change > 0.5 and p-value <0.05) were then matched to the human metabolome database^44^ using custom Matlab scripts^45^. For each tentative molecule that was identified, the biochemical pathways in which that molecule was involved were then identified.

Pathways were then reported in descending order of number of unique discriminating *m/z* that contained a molecule for that pathway (**Table S1**).

### Mass Spectrometry Imaging (MSI)

Frozen tissue samples were prepared for MSI as described previously^46^.

#### DESI-MSI

DESI-MSI was carried out using a Xevo G2-XS QToF instrument (Waters, UK) and a developmental sprayer incorporating a recessed capillary (Waters, UK). Solvent comprised of 95:5 methanol:water (Optima grade, Fisher Scientific, UK) was used. The DESI sprayer was operated at 0.8 kV, 4 bar Nitrogen gas pressure, 50 V cone, 100 °C ion block temperature, room temperature inlet capillary, sprayer angle of 80 degrees, 2 mm distance to the inlet capillary, 2mm distance from sprayer nozzle to sample. Prior to data acquisition DESI signal intensity was optimised using black permanent marker to reach ion intensity for *m/z* 666 of > 1e^6^ and on bovine brain homogenate tissue of > 1e^4^ for basepeak in lipid *m/z* region (700-900). Mass calibration was carried out daily or prior to each sample analysed, whichever was more frequent, for *m/*z range 50-1200 using mass spectra derived from a polylactic acid coated glass slide. Data were also acquired with this mass range for pixel size 75 × 75 µm at a scan rate of 2 pixels/second. DESI MS/MS was undertaken using the same conditions and instrument. The instrument was operated in the MS/MS mode with RF voltage set to 7.4. For identification of the precursor ion that was assigned as glucose (**Figure S2**) the collision energy used was 10.

Volcano plot produced from MSI data as shown in Figure S4A was produced from tumour sub-regions of MSI data defined by tSNE and k-means clustered ROIs which were observed to correlate with tumour regions in the corresponding H&E stained tissue sections from the same tissues. Pixel data were RMS normalised, zeros removed, and two sided t-tests were performed in Python (ttest_ind, scipy.stats package). Fold changes (FC) were calculated from average intensity values and plotted against p-value where 1.5 and 0.05, respectively, were considered as thresholds in the displayed figures.

#### MALDI-MSI

MALDI MSI was carried out using a Synapt G2-Si QToF instrument (Waters, UK) fitted with a uMALDI ion source^47^. The samples were coated with 9-AA (Merck Life Science, Dorset, UK) at 10 mg/mL in 80:20 ethanol:water by TM Sprayer (HTX Technologies, NC, USA) with a temperature of 65 °C, flow 0.06 mL/min, nozzle velocity 1200 mm/s, 4 passes, 3 mL track spacing, CC pattern alternation. Instrument was calibrated with Red Phosphorus to an RMS mass accuracy of < 1 ppm and instrument method was set as follows: pixel size of 45 × 45 µm, sensitivity mode, negative ion mode, *m/z* range 50-1200, laser repetition rate 2500 Hz, scan time 0.5 s.

#### DESI and MALDI MSI t-SNE Data Analysis

Data were converted from proprietary .RAW format to imzML using ProteoWizard^40^ and imzML Converter^41^; and analysed using SpectralAnalysis^42^ and in-house developed scripts in MatLab (2017a). Next, the top 4000 peaks of each data set were used to create a datacube and perform spatial segmentation by k-means clustering, with the similarity metric set to “cosine” and the number of clusters set to 4. Tissue and background were manually labelled by comparing the optical images with the spatial segmentation results. The total spectrum for the tissue was created by summing all pixels belonging to the clusters categorized as such. Following removal of background pixels, t-SNE was performed on the tissue-only pixels using the Matlab function “tsne”, the data were reduced to three dimensions, and the spectral similarity metric used was correlation. The remaining t-SNE parameters e.g. perplexity were set to their MATLAB default values. K-means cluster value (k) was determined using the elbow method^48^ unless otherwise stated. Data representation shown in Figure 2E was produced from background subtracted data with subsequent k=3 k-means clustering applied.

#### LESA MS/MS

LESA CID MS/MS was carried out using a SYNAPT G2-Si QToF (Waters, UK) and the LESA Advion TriVersa NanoMate was used as the ionisation source. A surface extraction solvent of 95:5 methanol:water was used for all experiments. LESA parameters used for analysis: solvent volume 4 µL, solvent depth 1 mm up from the bottom of the reservoir, dispensed 2 µL, delayed 2 sec post dispense, aspirated 3.5 µL, repeated mix 2 times, delayed 2 sec post aspirate. Negative ion nano-ESI was used for the MS/MS analysis with spraying parameters of 0.3 psi gas pressure and 1.4 kV for applied voltage. The SYNAPT G2-Si was operated in negative ion MS/MS mode for precursor ion at *m/z* 464. A range of CEs were used with the data averaged.

### Bioinformatic analysis

To study the gene expression of *AHCY* across the different consensus molecular subtypes (CMS) of CRC (Figure 3F), patients with colonic adenocarcinoma from the TCGA database were assigned to CMS classes using the CMSclassifier R package as described in Guinney *et al*.^15^. Differential gene expression for each CMS group relative to all others was then performed using the R package limma (version 3.48.3) as follows. Linear models were fitted per gene using the lmFit() function, then statistics were calculated using the eBayes() function before being extracted and multiple testing correction performed using the topTable() function, all with default settings.

### Data visualization and statistical analysis

Unless mentioned differently, data were plotted and analysed using GraphPad Prism 9.0.

### Data availability

All data relevant to this study are available from the corresponding author at reasonable request.

### Code availability statement

Processing scripts used for data mining and presentation will be uploaded to Github repository, which will be made public on publication of this manuscript.

## Acknowledgments

We thank all members of the Grand Challenge Rosetta consortium, Eyal Gottlieb, John R.P. Knight, Jan Balzarini and Catherine Winchester for discussion of the data and manuscript. We are grateful to the CRUK Beatson Institute Central Services, Molecular Technology Service, Histology Service, the Transgenic Technology Laboratory, and the Biological Services Unit for technical support. O.J.S. was supported by grants from CRUK (A21139, A25045, A17196, and A31287). J.V.V., A.K.N, D.M.G., R.T.S., C.J.N., A.D., L.Z., E.E., A.N., A.G., T.M., C.F., A.B., B.Y., and J.B. were supported by the CRUK Rosetta Grand Challenge grant (A25045 to J.B. and O.J.S.). D. S., C.N., R.A.R, and A.D.C. were supported by the CRUK Beatson Institute core grant (A17196 to O.J.S.). N.V. was supported by the Wellcome Trust (201487 to O.J.S.). K.G. was supported by the CRUK Grand Challenge Specificancer Grand Challenge Consortium (A29055 to O.J.S.). T.R.M.L. was funded by CRUK Accelerator Programme Award (A26825 to O.J.S.). O.D.K.M. was funded by a CRUK Career Development Fellowship (C53309/A19702). M.G. was funded by the University of Glasgow. This work was supported by the Francis Crick Institute, which receives its core funding from Cancer Research UK (FC001223), the UK Medical Research Council (FC001223) and the Wellcome Trust (FC001223, FC0010060); and by the CRUK Grand Challenge Award 2015 C57633/A25043.

## Author Contributions

J.V.V., A.C., M.Y., J.B, and O.J.S. designed the research. J.V.V., A.K.N, L.Z., M.G., and K.S. performed experiments and analysed data. R.T.S., C.J.N., A.D., E.E., A.G., A.N., T.M., A.B., and B.Y. performed REIMS and MSI, and analysed the data. D.S. and E.S. performed LC-MS, and D.S performed untargeted data analysis. C.A.F, T.R.M.L, N.V., D.M.G. and R.R. performed *in vivo* experiments. C.N. performed IHC and ISH. W.C. performed RNA sequencing. K.G. analysed RNA expression data. V.S.W.L. provided support with *Apc*^Min^ studies. O.D.K.M. provided help with stable isotope labelling studies. R.J.A.G, S.T.B., V.W, Y.X, and Z.T. provided advice on MSI and REIMS. J.V.V., A.C., and O.J.S. wrote the paper. All authors read the manuscript and provided critical comments.

## Tables

**Table S1. Pathway analysis of ions detected by REIMS, and found to discriminate between genotypes in epithelial extracts of WT, KRAS, APC, APC KRAS and APC KRAS PTEN mice.**

**Table S2. Tentative identification of ions detected by REIMS, and found to discriminate between genotypes in epithelial extracts of WT, KRAS, APC, APC KRAS and APC KRAS PTEN mice.**

**Table S3. List of ions (*m/z* values) used to construct volcano plot (Figure S4A) showing metabolic differences between tumour epithelial regions of APC and APC KRAS tumours analysed by DESI MSI (negative polarity)**

**Table S4: List of molecules (neutral mass) used to construct volcano plot (Figure S4C) showing metabolic differences between bulk tissue extracts from APC and APC KRAS tumours analysed by LC-MS (polarity switching)**

---

**Figure S1.**
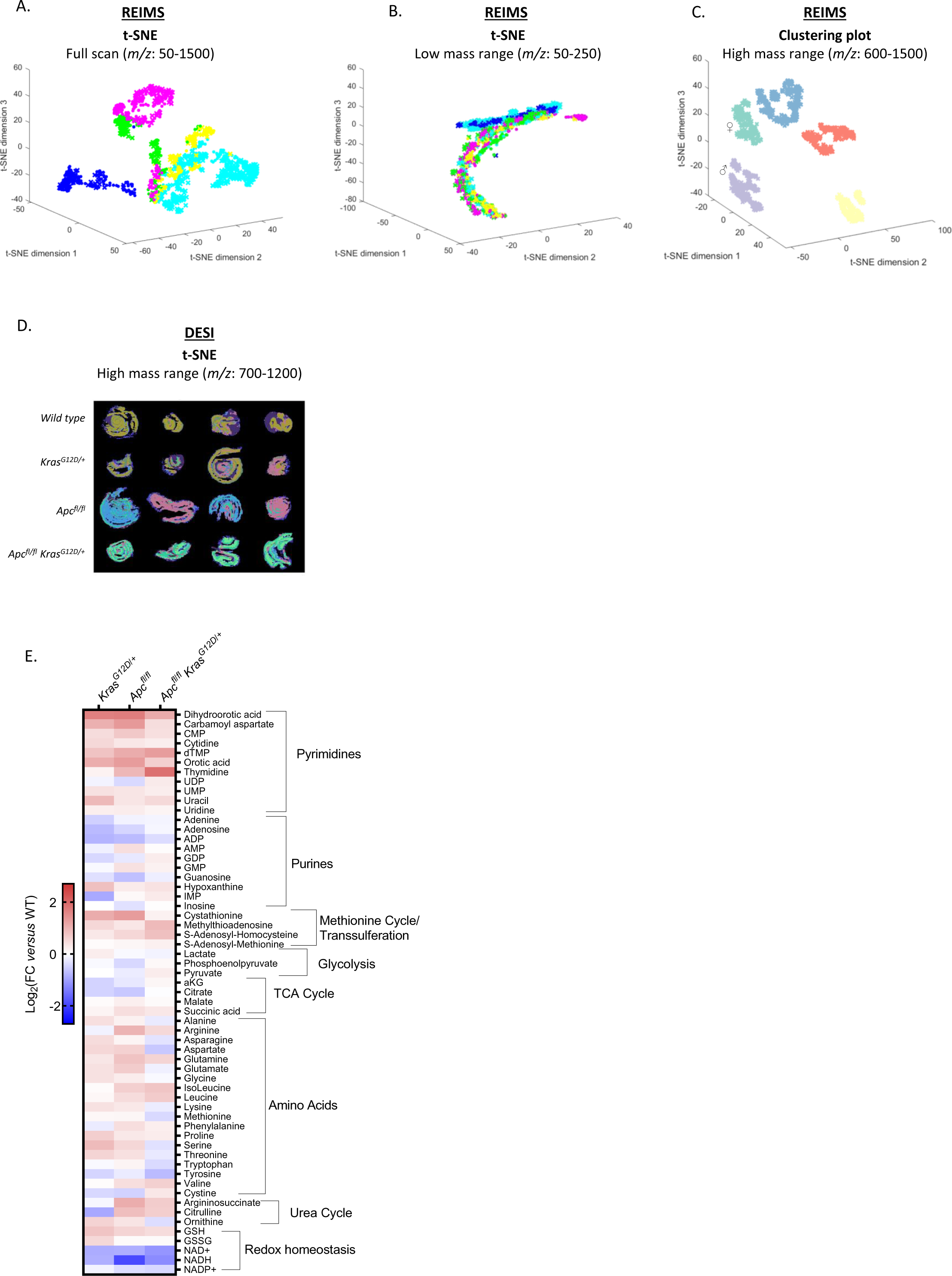
Metabolic profiling of intestinal tissues by REIMS, DESI-MSI and LC-MS. (A) Multivariate analysis of REIMS data showing segmentation when focusing analysis on the full acquired mass range (*m/z*=50-1500), or (B) low mass range (*m/z*: 50-250). Colour coding: yellow = WT; green = KRAS; cyan = APC; pink = APC KRAS; blue = APC KRAS PTEN. Each dot corresponds to a single mass spectrum acquired using the REIMS forceps. Data acquired from WT (n=4), KRAS (n=4) APC (n=11), APC KRAS (n=4), and APC KRAS PTEN (n=5) mice. (C) Clustering plot of REIMS data (*m/z*: 600-1500) showing genotype-dependent clustering. ♂/♀ indicate male/female dominant APC clusters. (D) t-SNE of DESI-MSI data acquired from longitudinal sections of the rolled up small intestine of WT, KRAS, APC, and APC KRAS mice focusing analysis on ions within mass range *m/z*: 700-1200 (n=4 per genotype, each image is rolled small intestine from one individual mouse). (E) Targeted LC-MS analysis of polar extracts of colonic tissues from KRAS, APC, and APC KRAS mice compared with WT mice. Heatmap constructed based on fold change between the averages of each experimental group (n=3).

**Figure S2.**
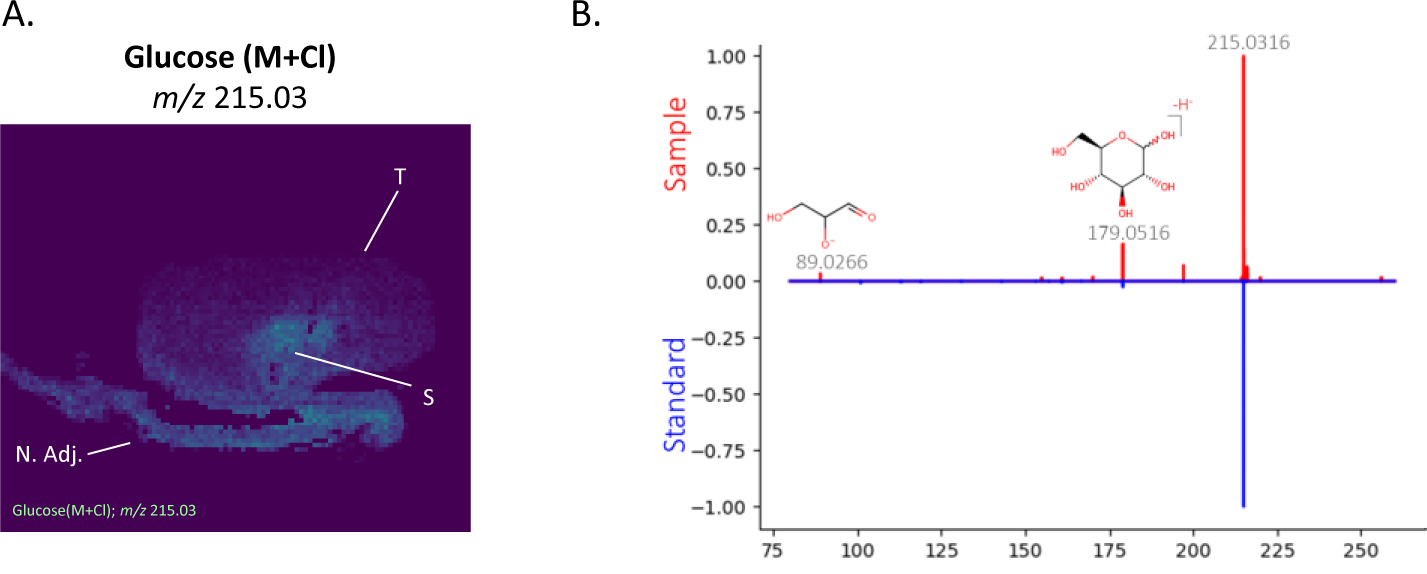
Decreased glucose abundance in tumour epithelium. (A) DESI-MSI showed decreased abundance of an ion with *m/z* 215.03 in the tumour epithelial compartment of APC tumours, and APC KRAS tumours (data not shown). [N. Adj.: normal adjacent tissue; S: stroma; T: tumour tissue]. Database search (https://hmdb.ca) suggested that this could be assigned as glucose [M+Cl]^-^. A standard solution of glucose was mixed with chlorinated dopamine to obtain chlorinated glucose for comparison. (B) DESI tandem mass spectrometry showed that the fragmentation of the ion of interest produced ions matched to the fragmentation pattern of the chlorinated standard, further supporting the ID of this metabolite of interest. All highlighted *m/z* of fragments and precursor ions on the mass spectra are present in both standard and sample.

**Figure S3.**
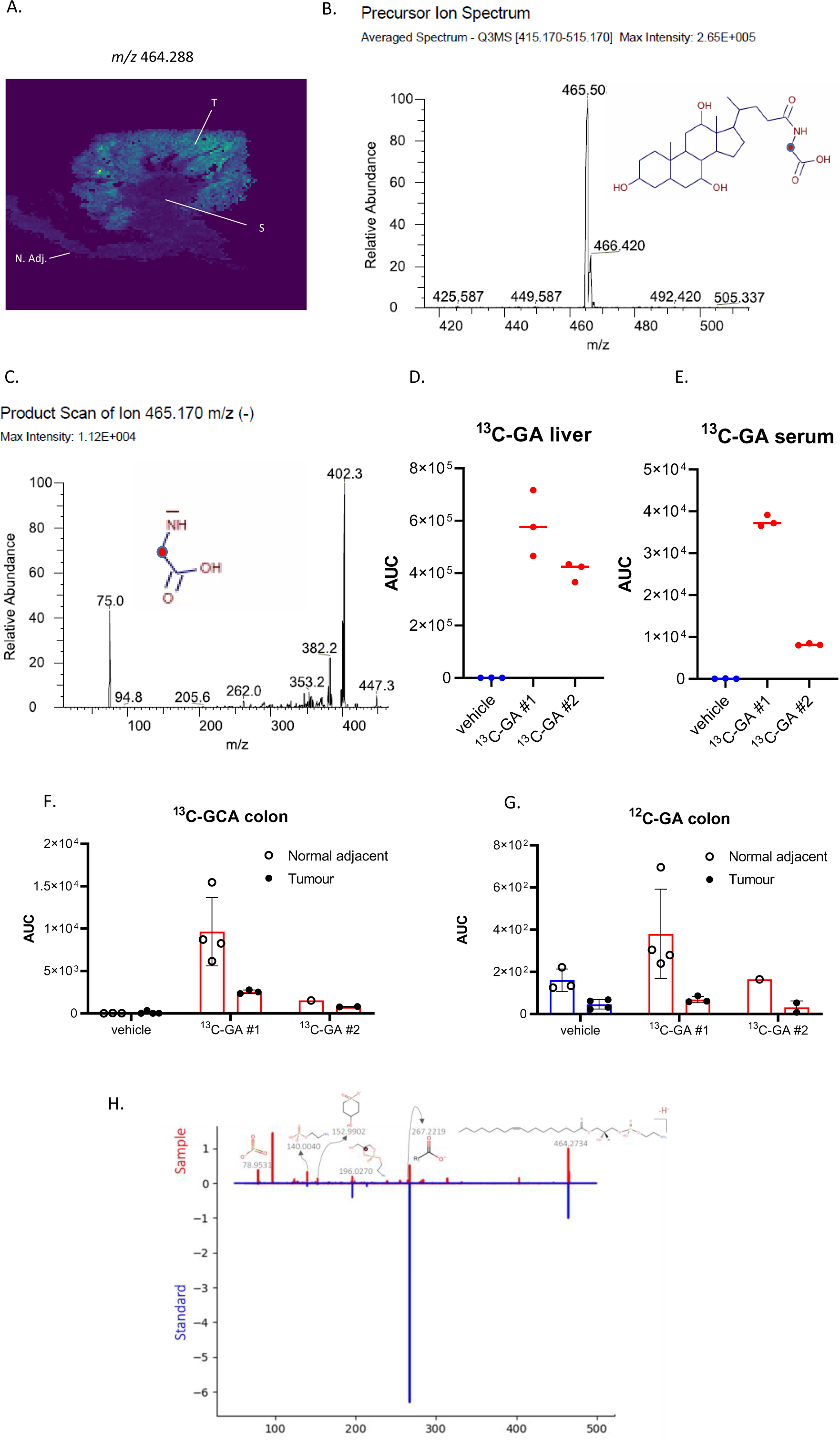
Tumour epithelial specific accumulation of Lyso-PE 17:1. (A) MALDI-MSI showed specific accumulation of an ion with *m/z* 464.288 in the tumour epithelial compartment of APC tumours [and APC KRAS tumours (data not shown)]. [N. Adj.: normal adjacent tissue; S: stroma; T: tumour tissue]. The two top tentative parent ion identifications (i.e. glycocholic acid and LysoPE 17:1) obtained from a publicly available database (https://hmdb.ca) were experimentally validated. (B) A ^13^C-glycocholic acid (^13^C-GA) standard solution was infused to detect the parent ion (red dot indicates ^13^C label) using Triple Quadrupole Mass Spectrometry, and (C) an MRM method was developed to study the abundance of ^13^C-GA in tissues via its a diagnostic ion of 75 Da (red dot indicates ^13^C label). Tumour-bearing APC mice were administered ^13^C-GA (75 mg/kg p.o.; n=2) or vehicle (HPMC/Tween:DMSO; v:v 90:10; n=1). After 8.5 hours, serum, liver and distal colon were harvested and processed for LC-MS. ^13^C-GA was detected in (D) liver, (E) serum, (F) normal adjacent colonic tissue and tumour tissues. No increased abundance of (F) ^13^C-GA or (G) ^12^C-GA was observed in tumour tissues compared with normal adjacent colonic tissue. Panels D-G show the mean of metabolic extractions of the multiple tissue fragments for each mouse, each dot represents data obtained from a single tissue fragment, or serum extract. Liquid extraction surface analysis tandem mass spectrometry was applied to study fragmentation of the *m/z* 464.288 precursor ion, which showed matched ions to the predicted fragmentation pattern of LysoPE 17:1, as well as to the (H) fragmentation of a commercially available standard further supporting the ID of this metabolite of interest. All highlighted *m/z* of fragments and precursor ions on the mass spectra are present in both standard and sample.

**Figure S4.**
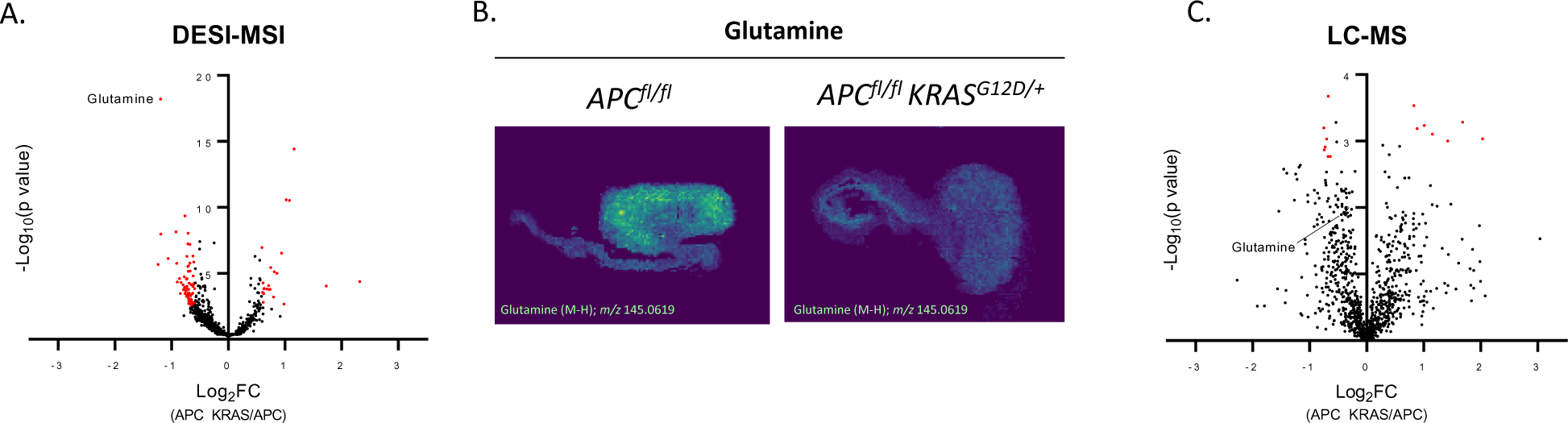
Comparative untargeted metabolomics of APC and APC KRAS tumours by DESI-MSI and LC-MS. (A) Volcano plot showing metabolic differences between tumour epithelial regions of APC (n=2 mice) and APC KRAS (n=2 mice) colon tumours as analysed by DESI-MSI. Red dots: FC ≥ 1.5 and significant after Benjamini-Hochberg FDR correction (q=0.1). (B) Representative images of glutamine abundance in APC and APC KRAS colon tumours as analysed by DESI-MSI. (C) Volcano plot showing metabolic differences between bulk tumour tissue extracts of APC (n=5 mice) and APC KRAS (n=8 mice as analysed by untargeted LC-MS. Red dots: FC ≥ 1.5 and significant after Benjamini-Hochberg FDR correction (q=0.1).

**Figure S5.**
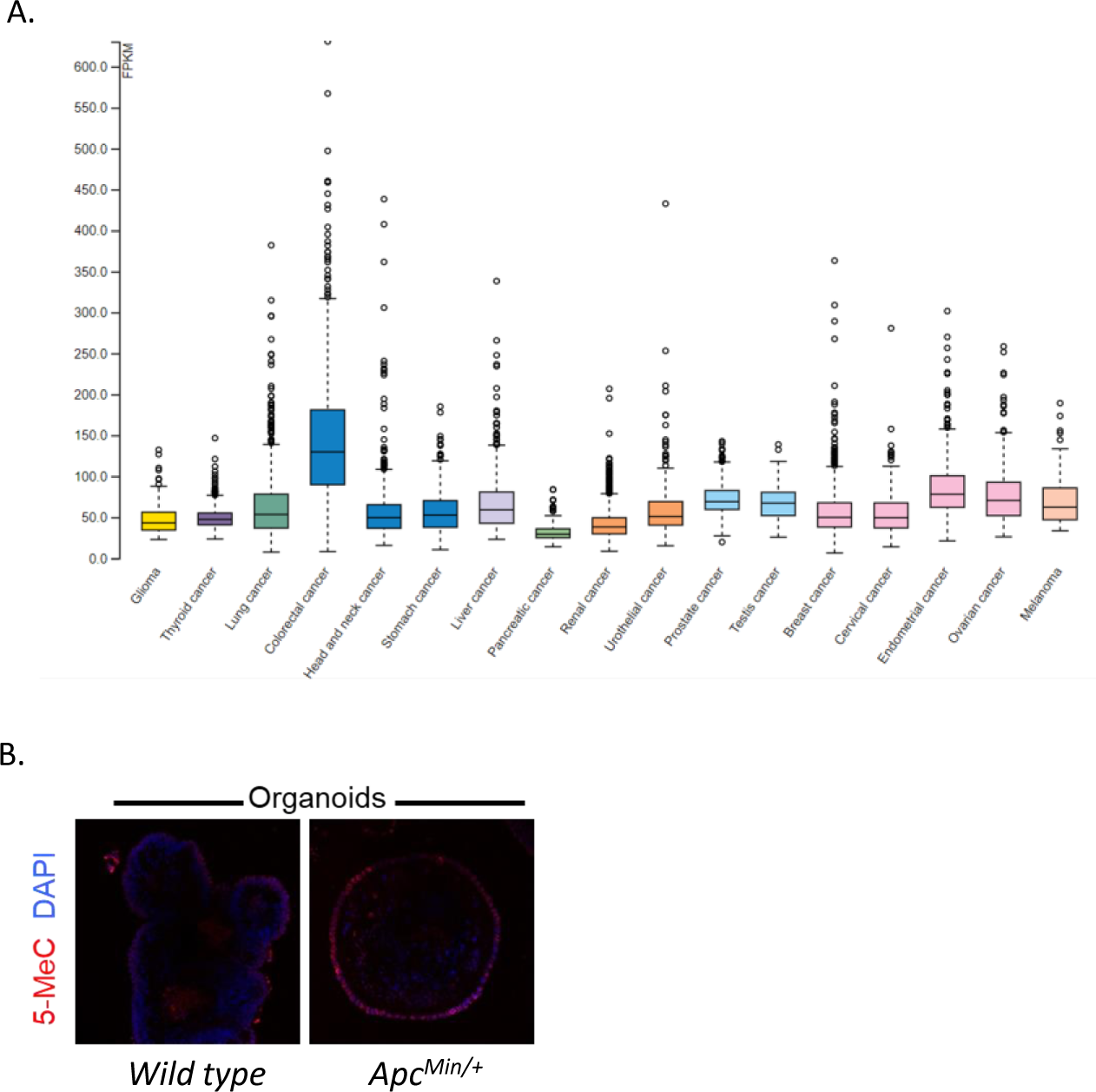
*AHCY* expression in different cancers, and 5-methylcytosine abundance in intestinal organoids. (A) Pan-cancer analysis showing *AHCY* gene expression in 17 cancers as analysed by RNA-Seq. Data reported as FPKM (number fragments per kilobase of exon per million reads), generated by The Cancer Genome Atlas (TCGA). Plot obtained from https://www.proteinatlas.org/. (B) IF showing abundance of 5-methylcytosine in intestinal organoids isolated from WT or *APC^Min/+^* mice.

**Figure S6.**
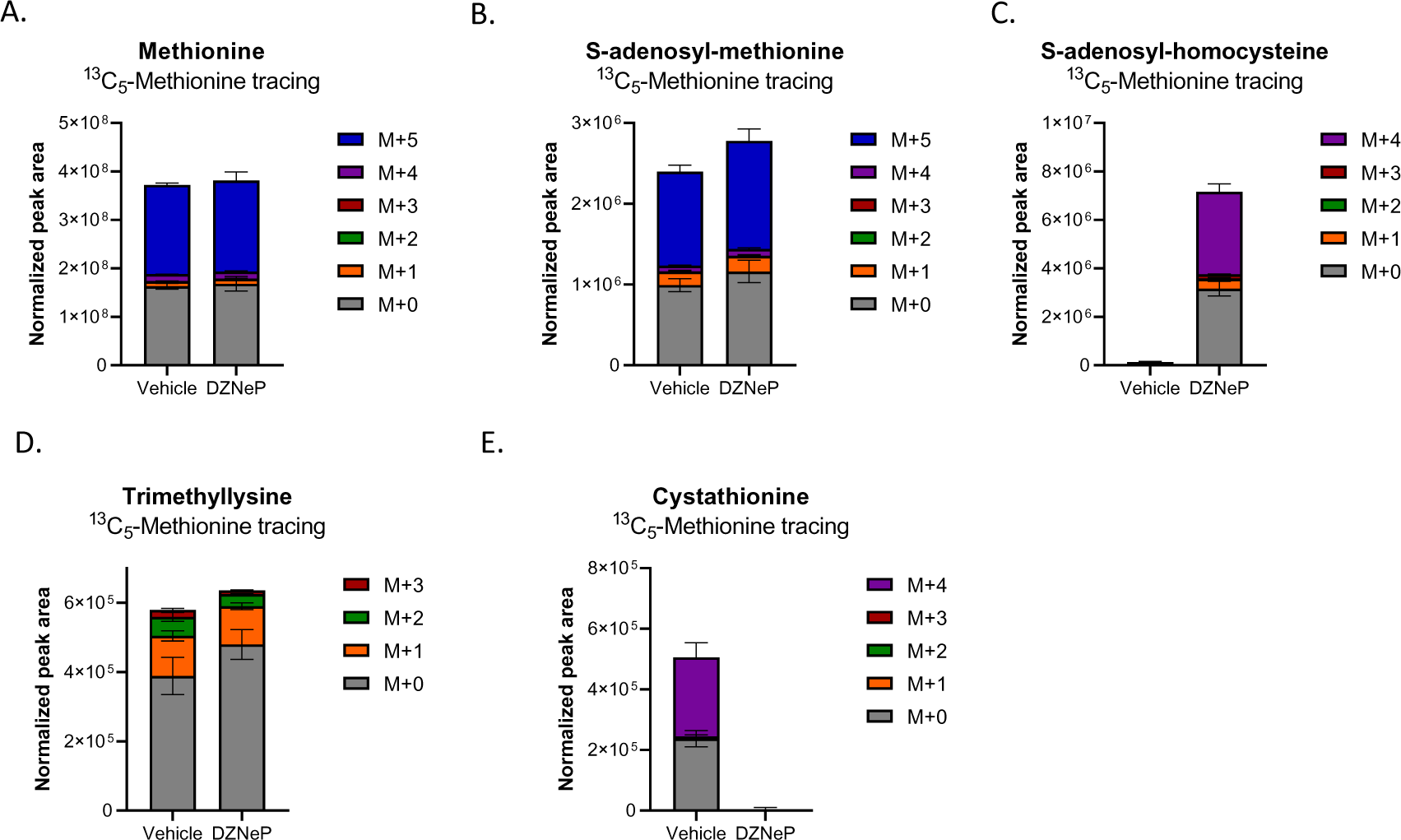
Methionine tracing in APC organoids treated with vehicle or DZNeP. Plots accompanying Figure 4(D-H) showing the abundance of all isotopologues for (A) methionine, SAM, (C) SAH, (D) trimethyllysine, and (E) cystathionine in APC organoids (+/- DZNeP 1 μM) cultured in the presence of ^13^C_5_-methionine. Data from a representative experiment performed twice, with 4 technical replicates each. Mean ± SD.

**Figure S7.**
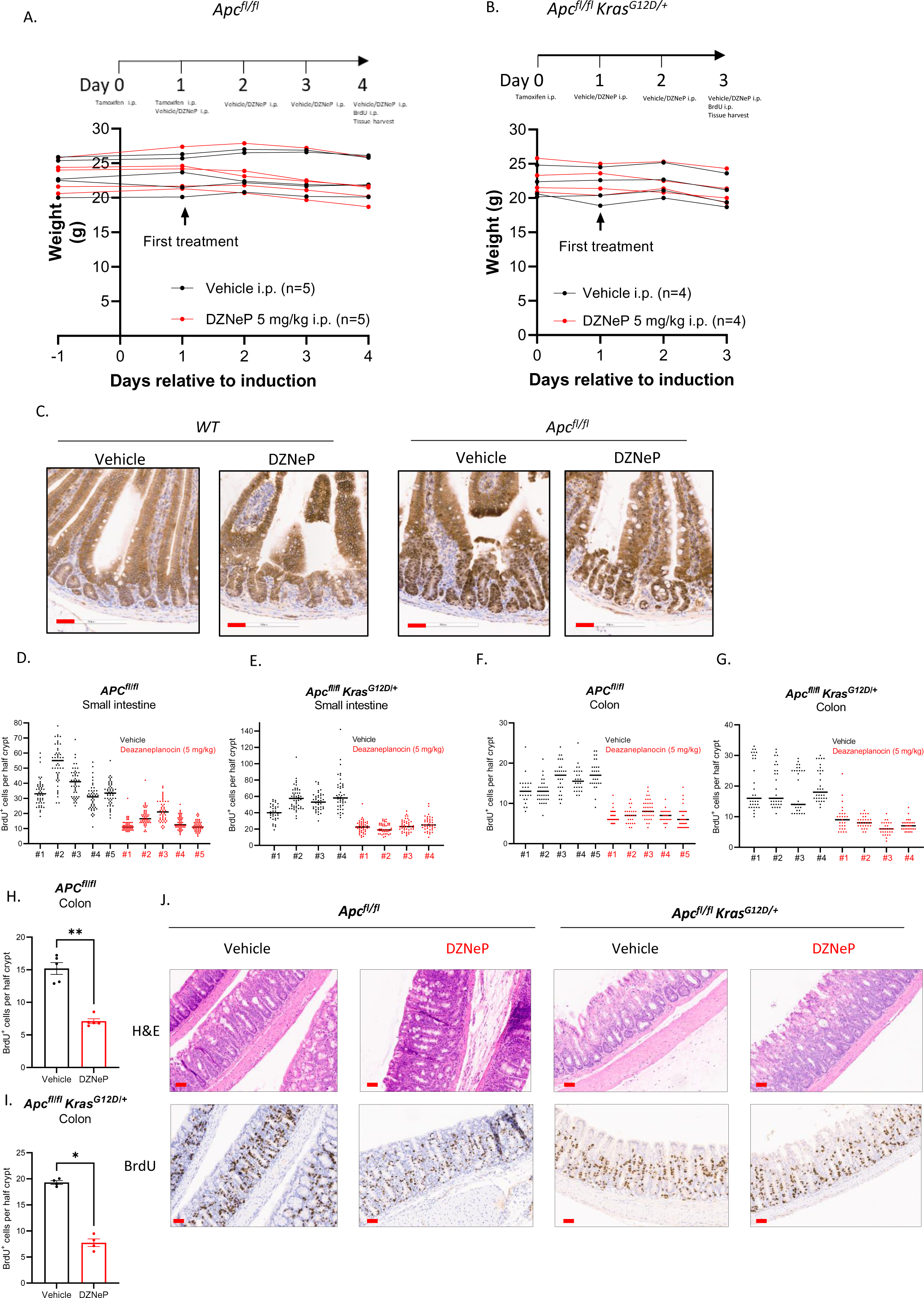
Effects of DZNEP on β-catenin localization and intestinal proliferation in APC and APC KRAS mice. (A,B) Schematic representation of experiments performed to test the effect of DZNeP in APC and APC KRAS mice, and effect of treatment on body weight. (C) Representative images of IHC for β-catenin in the small intestine of WT (n=3) and APC (n=5) animals treated with vehicle or DZNeP (5 mg/kg). Scale bars (red): 50 μm. (D-G) Quantification of IHC for BrdU incorporation in the (D,E) small intestine and (F,G) colon of APC and APC KRAS mice treated with vehicle or DZNeP (5 mg/kg). [n=5 (APC) or n=4 (APC KRAS) mice per experimental arm; Individual data shown for each mouse. Each dot represents the number of BrdU positive cells counted per half crypt. (H,I) Quantification of IHC for BrdU in the colon of APC and APC KRAS mice treated with vehicle or DZNeP (5 mg/kg). [n=5 (APC) or n=4 (APC KRAS) mice per experimental arm; Mean ± SEM, Each dot represents the average number of BrdU positive cells per half crypt for each mouse]. Asterisks refer to p-values obtained from 1-tailed unpaired Mann-Whitney tests (*: p<0.05; **: p<0.01). (J) Representative images of H&E staining and IHC for BrdU on colon sections of APC (n=5) and APC KRAS (n=4) mice treated with vehicle or DZNeP (5 mg/kg). Scale bars: 50 μm.

**Figure S8.**
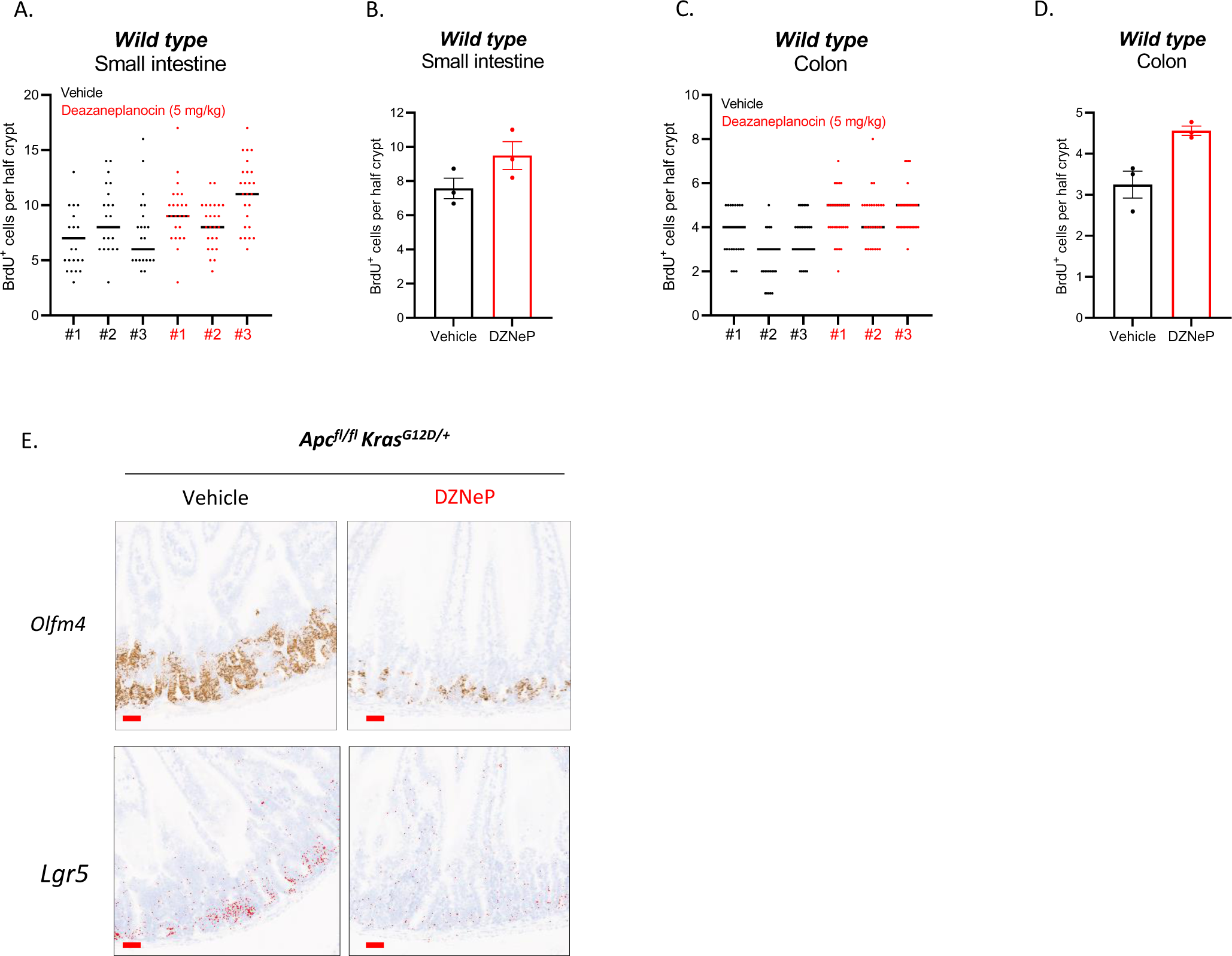
DZNeP treatment in WT mice and effect of DZNeP on stem cell markers in APC KRAS mice. Quantification of IHC for BrdU incorporation in the (A,B) small intestine and (C,D) colon of WT mice treated with vehicle or DZNeP (5 mg/kg). (n=3 mice per experimental arm; panels A,C: each dot represents the number of BrdU positive cells counted per half crypt; panels B,D: Mean ± SEM, each dot represents the average number of BrdU positive cells per mouse. (E) Representative images of ISH for *Olfm4* and *Lgr5* expression in the small intestine of APC KRAS mice (n=4) treated with vehicle or DZNeP (5 mg/kg). Scale bars: 50 μm.

